# Causal interactions from proteomic profiles: molecular data meets pathway knowledge

**DOI:** 10.1101/258855

**Authors:** Özgün Babur, Augustin Luna, Anil Korkut, Funda Durupinar, Metin Can Siper, Ugur Dogrusoz, Joseph E. Aslan, Chris Sander, Emek Demir

## Abstract

Measurement of changes in protein levels and in post-translational modifications, such as phosphorylation, can be highly informative about the phenotypic consequences of genetic differences or about the dynamics of cellular processes. Typically, such proteomic profiles are interpreted intuitively or by simple correlation analysis. Here, we present a computational method to generate causal explanations for proteomic profiles using prior mechanistic knowledge in the literature, as recorded in cellular pathway maps. To demonstrate its potential, we use this method to analyze the cascading events after EGF stimulation of a cell line, to discover new pathways in platelet activation, to identify influential regulators of oncoproteins in breast cancer, to describe signaling characteristics in predefined subtypes of ovarian and breast cancers, and to highlight which pathway relations are most frequently activated across 32 cancer types. Causal pathway analysis, that combines molecular profiles with prior biological knowledge captured in computational form, may become a powerful discovery tool as the amount and quality of cellular profiling rapidly expands. The method is freely available at http://causalpath.org.

## Introduction

Molecular pathways are models grounded in biochemistry that explain how perturbations are related to observed molecular and cellular responses. Due to the ubiquitous presence of irreversible reactions, they are often conveyed in terms of cause and effect. As early as 1950s, large pathway models were compiled from sets of carefully designed low throughput controlled experiments in a piecemeal fashion. They proved themselves to be powerful models that can be used to hypothesize about the effects of previously unobserved perturbations.

Advancement of -omic technologies opened a multi-layer and high-throughput window into the cell, presenting an opportunity—and a great challenge—for inferring large scale, causal pathway models. A diverse set of algorithms have been developed to solve this problem on proteomic and other associated molecular data^1–3^. They have different strengths and weaknesses, but they typically require many perturbations and time-resolved profiling, which is impractical for most research studies. Not testing every possible combination of perturbations leads to less confident pathway inference because different result models would equally fit to the data. A remedy is to use “prior information” in the algorithms, which roughly means, whenever multiple solutions exist, to choose the one that is supported by a known pathway relation that was discovered in another context. The hidden assumption in this approach is that a pathway fragment can be applicable to multiple contexts, and their applicability is more likely if it is also supported by profiling data. We and others have previously shown that using prior information increases predictive accuracy of pathway inference methods^4,5^.

The high-throughput aspect of the current -omic technologies only applies to the number of measured molecules, and not to the perturbations or time-resolving. Each perturbation has to be carefully designed in a low throughput setting, increasing costs linearly. The vast majority of research projects use proteomics with minimal perturbations such as before/after a drug treatment, or no perturbations at all such as profiling of a disease cohort. As molecular datasets get poorer in perturbations, they have to depend more and more on priors for pathway inference. But this scheme has one problem: priors used in pathway inference methods are generally simple networks such as protein-protein interactions—they do not inherently suggest any causal relationship between measured molecules. To extend pathway inference to more common proteomic datasets using priors, we need high-confidence priors that contain causality within. In this study, we propose a method for generating causal priors from mechanistic pathway relationships curated from scientific literature, then we use them for the pathway analysis of publicly available perturbation-poor proteomic datasets. Our approach reduces the pathway inference task into identifying the causal priors that can explain correlations in a given molecular dataset, which we distinctly name as pathway extraction. Pathway extraction, of course, can not fully cover for the lack of rigorous perturbations, and it is limited by previously identified pathway fragments in other contexts, but it generates extremely useful and falsifiable hypotheses that are unavailable otherwise.

Our method—CausalPath—uses curated mechanistic human pathways from multiple resources that are integrated into the Pathway Commons database^6^, detects the causal links in the pathways between measurable molecular features using a graphical pattern search framework, and identifies the subset of the causal links that can explain correlated changes in a given set of proteomic and other molecular profiles. We render these explanations as an intuitive simplified network, but also link to the relevant detailed models and the related literature to establish a powerful analysis platform (Fig. 1). This approach, in essence, mimics the literature search of a biologist for relationships that explain their data. Automating this task enables asking multiple interesting questions that would be too time-consuming to do through manual analysis. Since this process systematically considers hundreds of thousands curated mechanisms, it also is more comprehensive, unbiased, and more consistent in terms of the generated hypotheses.

**Figure 1.**
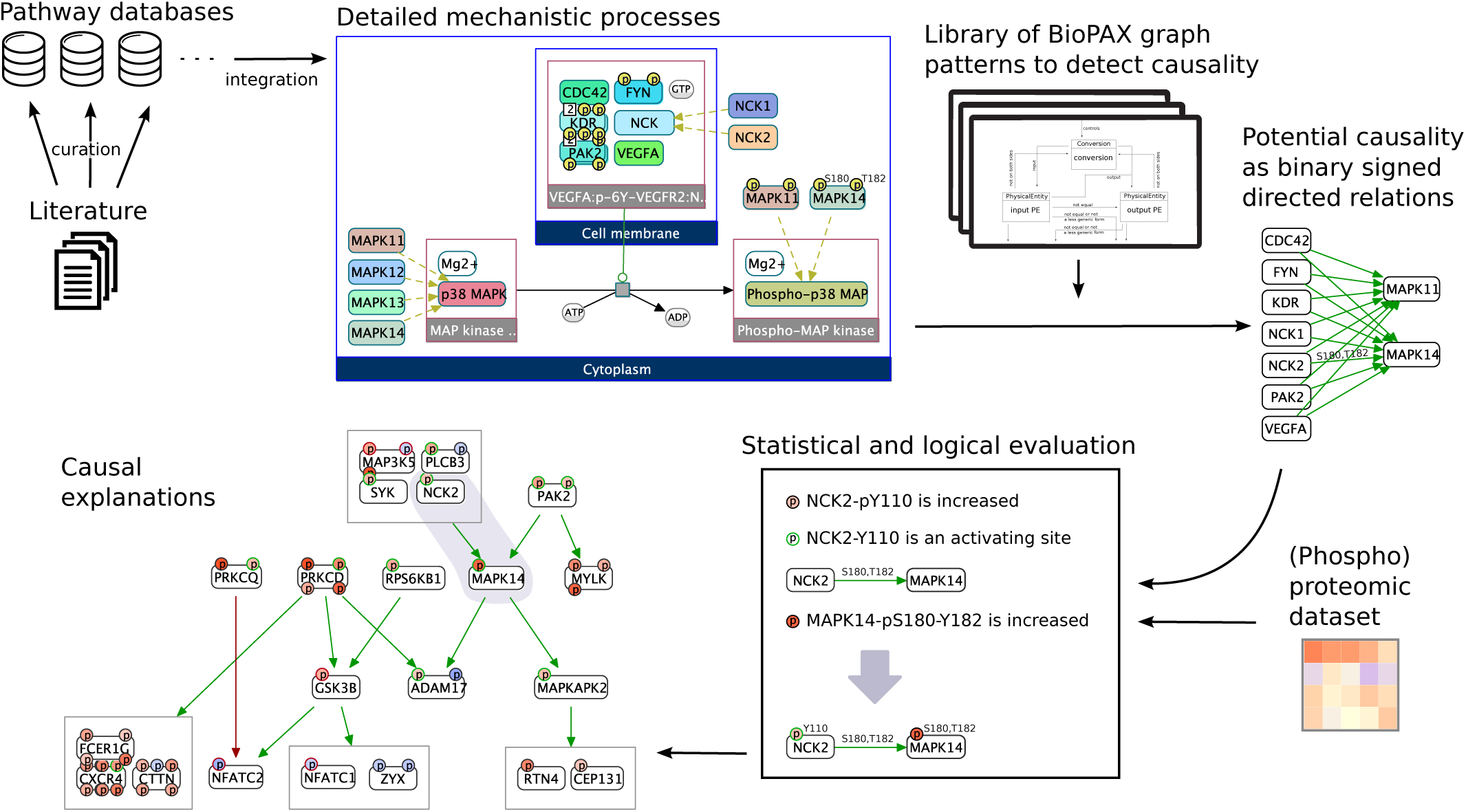
Design of CausalPath. A walk-through over components showing how a specific result relation (NCK2 → MAPK14) was generated for the platelet activation study. Information in scientific literature gets curated into pathway databases, which we integrate into Pathway Commons as detailed mechanistic processes. We detect structural patterns in these processes that can causally link two measurable protein features to generate binary causal priors. The statistical and logical evaluation step detects which causal priors have support in the proteomic dataset. The final logical network shows the 20 relations in platelet activation analysis results. For a description of graph notation please see Figure 2C. We omit phosphorylation site locations while rendering the result network for complexity management. These are revealed to the user upon clicking on nodes or edges in our interactive tools.

We tested this framework on an EGF stimulation dataset, detected EGFR activation with its signaling downstream to MAPKs, and a feedback inhibition on EGFR itself. We analyzed platelet activation and detected a plausible link from MAPK signaling to apoptotic signaling, which we validated by perturbing MAPK proteins. We analyzed ovarian cancer and breast cancer mass spectrometry datasets from two CPTAC projects, focusing on general and subtype specific signaling, as well as detecting regulatory relations over well-known cancer proteins. We analyzed RPPA datasets from 32 TCGA projects and found that AKT signaling is the most frequently varied signaling among patients in the same study.

## Results

### Design and properties of CausalPath

CausalPath work flow has two main components: (i) generation of causal priors from pathway databases—performed once and reused in multiple analyses, (ii) matching causal priors with supporting correlated changes in the analyzed data—performed for every analysis. We define a *causal prior* as a set of prior knowledge that together suggest a possible causal link between two measurable molecular features.

Although simple kinase-substrate databases and transcription factor-target databases exist they are limited for a comprehensive causal reasoning. Many kinases, phosphatases and transcription factors function as parts of molecular complexes curated by the pathway databases that support detailed mechanistic models. Such models can use multi-level nesting, generalizations such as homologies, and non-trivial mechanisms such as the use of small molecule secondary messengers. To detect structures that imply causal relationships between proteins in the Pathway Commons database, we used the BioPAX-pattern software, and manually curated 12 graphical patterns (described in Supplementary Doc). Searching for these patterns in Pathway Commons generated 20,032 phosphorylations, 2,767 dephosphorylations, 4,921 expression upregulations, and 811 expression downregulations as signed and directed binary relations. To increase coverage, we added relations from several other databases (PhosphoNetworks^7^, IPTMNet^8^, TRRUST^8^ and TFactS^9^) which are not in Pathway Commons, and increased our relationships by 24433, 2767, 9060 and 2961, respectively.

We define a causal conjecture (CC) as a pairing of a causal prior with supporting measurements in the molecular dataset that together declare that “one molecular change is the cause of another molecular change”. As an example, consider a platelet activation study that detects a set of proteomic changes. The chain of items below forms a causal conjecture:

1. NCK2-pY110 peptide level is increased in response to platelet activation.
2. Y110 is an activating phosphorylation site of NCK2.
3. NCK2 is part of a complex that can phosphorylate MAPK14 at S180 and Y182.
4. MAPK14-pS180-Y182 peptide level is increased in response to platelet activation.

Items 1 and 4 are direct observations from proteomic profiles and they are observed within the platelet activation context, and items 2 and 3 are the knowledge fragments that construct the causal prior, demonstrated in different contexts, published and subsequently curated into pathway databases. The causal conjecture is that the increase in phospho-NCK (NCK2-pT110) caused an increase in its activity of phosphorylating MAPK14, hence an increase in MAPK14-pS180-T182 (Fig. 1) in the context of platelet activation. This is a well-defined, mechanistic and falsifiable conjecture that can be automatically generated and easily testable in the lab.

Alternative to the above controlled perturbation setting where we compare control and test samples, a causality network can be generated based on the correlations of measurements in an uncontrolled cohort—as is common in cancer biology. In that case, we replace items 1 and 4 with an observed correlation, e.g. “Measured peptide levels of NCK2-pT110 is positively correlated with the peptide levels of MAPK14-pS180-T182”, for a correlation-based causality hypothesis.

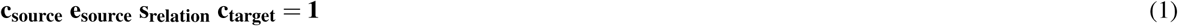

To formalize and generalize this example of causal conjecture detection, we can formulate it with a logical equation (Eq. 1), where c represents the change direction of the gene features, e represents the effect of the source feature on its activity, and s represents the sign of the pathway relation, where *c*, *e*, *s* ∈{−1,0,1}. Four terms in the equation correspond to the four items in the example, which collectively test if the data is consistent with a known causal interaction. The effect of source feature *e_source_* is 1 in the case of total protein and activating phosphorylation, and it is −1 in the case of inactivating phosphorylation. Relation sign, *s_relation_*, is 1 for phosphorylation and expression upregulation, and −1 for dephosphorylation and expression downregulation. Additionally, we limit the phosphorylational controls to the explanation of phosphoprotein changes, and limit the expressional controls to the explanation of total protein changes (or mRNA changes), which are not encoded in the equation. In the case of correlation-based causality, the term *c_source_c_target_* in Eq. 1 is replaced with the sign of the significant correlation between source and target features.

On top of the logic-based detection of causal interactions, we provide two types of statistical measurements that help to assess the confidence in the results. The first one checks if the proteomic data is in fact likely controlled by the known relations. If true, we expect to have a significantly high number of interactions in the results, which we test by data label randomization and recalculating the results multiple times. A second statistic checks if a protein on the network has significantly large number of downstream targets in the results. We test this for each protein using the same randomization procedure. Significant values provide additional evidence showing the data is not random, or a protein has visible influence on many targets, which consequently increases our confidence in the results.

We provide analyses for robustness and reproducibility of CausalPath results, as well as a survey of other methods related to pathway analysis for proteomic datasets in the Methods.

### Analysis of platelet activation

We recently used a preliminary version of CausalPath to analyze quantitative changes in the phosphoproteome of blood platelets stimulated with the purinergic receptor agonist ADP^10^. The aim of this study was to generate testable hypotheses regarding known and novel pathways that regulate platelet function. We identified 20 relations that causally explain correlated changes in 21 phosphopeptides (Fig. 1). One of these identified relations, phosphorylation of RTN4 (a BCL2L1 sequestration protein) at S107 by MAPKAPK2, was not previously described in the context of platelet activation, but suggested a link from MAPK14 (p38 MAPK) signaling to endoplasmic reticulum physiology and apoptotic signaling events that have central roles in the early events of platelet activation. To test this hypothesis put forth by CausalPath analysis, we examined the phosphorylation of RTN4 in activating platelets under control as well as MAPK14- and MAPKAPK2-inhibited conditions by Western blot and fluorescence microscopy. Ultimately, we verified that MAPK signaling specifically drives RTN4 phosphorylation in activating platelets^10^, demonstrating that CausalPath can generate novel, testable hypotheses for discovery.

### Analysis of EGF stimulation on EGFR Flp-In cells

We used a recent EGF stimulation time series phosphoproteomic dataset^11^ to see if CausalPath can recapitulate known biology, and to understand its limits. This dataset contains 8 time points, where the first time point (0 min) is unstimulated cells. We compared each of the other time points to the first time point to see how the cellular signaling evolves over time (Supplementary Animation 1). Since the data is phosphopeptide-only and does not contain any observable change on EGF itself, we included EGF activation as an hypothesis to the analysis. Consistent with our expectations, in the initial time points, CausalPath detects many EGFR targets and relates them to EGFR phosphorylation and activation. Both EGF and EGFR downstream are significantly enriched with changes that indicate their activation. At the 5th time point (16 min), we observe an inhibitory feedback phosphorylation of EGFR explainable by MAPK1 and MAPK3 activity, which follows with the disappearance of EGF signaling. All the networks up to the 5th time point are significant in size (*p <* 0.0001). What is missing in these graphs is an explanation for MAPK1/3 phosphorylation. It is known that EGF signaling can activate MAPK1/3 through several steps and multiple paths, but none was captured. One reason is that we are using CausalPath in the most strict configuration, forcing phosphorylation positions in the literature to exactly match the detected sites in the phosphoproteomic data. When we slightly relax this constraint by allowing 2 amino acids difference in site locations, we detect that SHC1 and GAB1 phosphorylations can causally link EGF stimulation to MAPK3 phosphorylation (Supplementary Animation 2). Our review of slightly shifted sites indicates that the site locations in the literature do not always map to the sequence of the canonical protein isoform provided by UniProt. For instance, one source of inconsistency is whether to count the initial methionine on the protein which is often cleaved. We are actively working on curation-correction tools for addressing this problem in the future. As a stop gap measure, CausalPath’s option to relax the site matching by 2 amino acids is useful for most applications.

### Analysis of CPTAC ovarian cancer dataset

Ovarian cancer is a form of gynecological cancer that was estimated to kill 14,240 women in USA in 2016^12^. High-grade serous ovarian cancer (HGSOC) is a molecular subtype which was subject to a TCGA project that investigated HGSOC with several kinds of molecular profiling on 489 patients^13^. HGSOC is often characterized by TP53 mutation (~95% of cases), low rates of other mutations, and extensive DNA copy number alterations. A recent CPTAC project performed proteomic and phosphoproteomic analysis on the 174 of the original TCGA ovarian cancer samples using mass spectrometry, providing measurements for 9,600 proteins from the 174 samples and 24,429 phosphosites from 6,769 phosphoproteins from 69 samples^14^.

Using CausalPath on this dataset, we generated explanations for the observed correlations in the measured peptide levels, using phosphorylation and expressional control pathway relations. The resulting phosphorylation network contains 116 relations and the expression network contains 249 relations when we use a 0.1 false discovery rate (FDR) threshold for correlations. Interestingly, while the size of the phosphorylation network is significantly large (*p <* 0.0001, calculated by data label randomization), we do not observe this for the expression network (*p* = 0.6482), suggesting that the data is shaped by known phosphorylations more than by known expressional controls. The most noticeable parts of the phosphorylation network include CDK1 and CDK2 downstream, MAPK1 and MAPK3 downstream, and several immune-related protein activities such as SRC family kinases, PRKCD, and PRKCQ (Fig. 2A).

**Figure 2.**
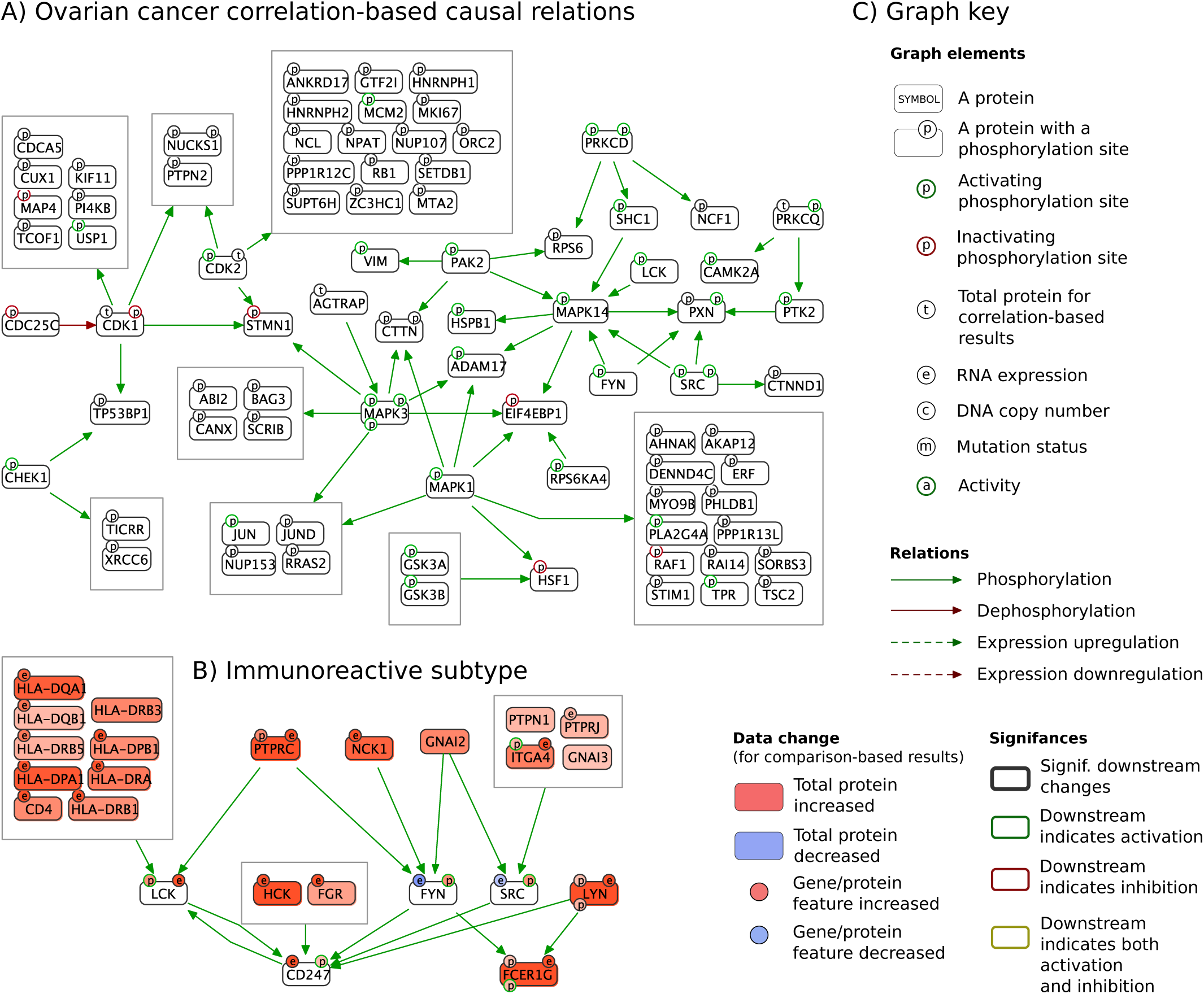
Results for CPTAC ovarian cancer. A) The largest connected component in correlation-based causality network. The complete network is in Supplementary Fig. 1. B) Immunoreactive subtype compared to all other samples, where we show RNA expression and DNA copy variation from corresponding TCGA datasets along with the CPTAC proteomic changes. C) Key for graph notation for causal explanations in all figures. See methods for a detailed explanation.

It is interesting that we obtain radically different significance values for expressional relations compared to phosphorylations. This could be either due to the low quality of expressional relations, higher number of confounding factors and/or due to the total protein measurements being a bad proxy for their RNA expression. To investigate this, we modified CausalPath to use TCGA RNAseq data instead of proteomic data for the target genes of expressional controls. We obtained 192 expressional relations that explain RNA measurements of 133 genes with proteomic changes of 89 transcription factors or their modulators (Supplementary Fig. 2). This time, the number of resulting relations is significantly large (*p* < 0.0001) suggesting that proteomic change is not a very good proxy for RNA expression. Additionally, the downstream changes of 5 transcription factors (STAT1, NFKB1, MCM6, CBFB and SPI1) are significantly large (0.1 FDR), promoting these factors as most influential for generating variance in ovarian cancer. It is counterintuitive to find less number of relations (192 versus 249) with higher significance (*p* < 0.0001 versus *p* = 0.6482), especially when the significance is based on the number of relations. One reason behind this can be the fact that only 106 of the 174 samples have RNAseq data available, hence, we have lower statistical power to detect correlations. But within relations that have significant correlations, they are ranked where we expect them to be by known causal relations.

The correlation-based causal network provides hypotheses for the signaling network parts that are differentially active across samples, but it does not tell which parts are activated together or whether they align with previously defined molecular subtypes. The original TCGA study on HGSOC samples identifies four molecular subtypes based on RNA expression, termed as immunoreactive, differentiated, proliferative, and mesenchymal^13^. To understand if there is a mapping from this network to the previously defined subtypes, we compared each subtype to all other samples but we were unable to generate results within 0.1 FDR threshold, probably due to the large proportion of missing values in the phosphoproteomic dataset combined with the loss of statistical power due to smaller cohort size for each subtype. Then we tried to constrain the search space with the neighborhoods of some of the genes with differential measurements, and relax the FDR threshold at the same time for further exploration. 6 SRC family kinases (SFKs) have proteomic evidence for activation in the immunoreactive subtype, hence, we limited the search to the neighborhood of SFKs (SRC, FYN, LYN, LCK, HCK, and FGR), set FDR threshold to 0.2 for phosphoproteomic data, and identified 27 relations (Fig. 2B). The network identifies several human leukocyte antigen (HLA) system (the major histocompatibility complex (MHC) in humans) proteins at SFK upstream, along with other genes regulating immune cell activation such as CD4, ITGA4, PTPRC, PTPRJ, PTPN1, and NCK1. On the network, we can track their signal going over SFKs to CD247 and FCER1G, immune response genes.

### Analysis of CPTAC breast cancer dataset

Another CPTAC project produced proteomic and phosphoproteomic profiles for 105 of the original TCGA breast cancer samples with mass spectrometry^15^. Unlike the ovarian cancer dataset, this dataset is rich in correlations, which can be explained by 2,324 phosphorylation and 1,497 expressional control relations. Similar to the ovarian cancer results, we detected the resulting phosphorylation network is significant in size (*p* < 0.0001) while the expression network is not (*p* = 0.8070), suggesting the known phosphorylation relations have a much higher impact on the proteomic correlations than known expressional relations. When we use TCGA RNAseq data instead of proteomic data for the targets of expressional relations, we detect 265 relations that explain RNA changes of 165 target genes by proteomic changes of 128 transcription factors or their modulators (Supplementary Fig. 4). This time, the size of the network is highly significant (*p* < 0.0001), with 7 transcription factors (STAT1, ESR1, MCM5, MCM6, GATA3, IL18 and S0X10) and MDC1 protein having correlated targets enriched in the results (0.1 FDR).

On the phosphorylation network, for complexity management and to focus on the most interesting part, we used a stricter FDR threshold of 0.001 and searched for regulators of known cancer proteins. We collected annotated “cancer genes” from OncoKB^16^ and COSMIC Cancer Gene Census^17^, and generated a subgraph with upstream neighbors of proteins related to those genes, which resulted in 205 relations with 98 cancer proteins controlled by 79 other proteins in breast cancer (Fig. 3A). Among 79 proteins, we detected that MAPK14 is the most influential one, controlling 11 cancer proteins, followed by CDK2 controlling 10 cancer proteins on the network. Targeting CDK2 and other CDKs is an ongoing effort in cancer therapy with a challenge of toxicity^18,19^. MAPK14 has conflicting reports on its anti- and pro-tumor activity^20^. This complexity and toxicity is expected given that these proteins exert control over too many important proteins, the effect of targeting them will likely depend on the current state of their downstream proteins in a patient. A better strategy may be to direct research towards personalized treatments where patients are profiled for the activity of known cancer proteins and the relevant regulators are targeted. In such a setting, the causal relations we identified would be very useful to decide on a target.

**Figure 3.**
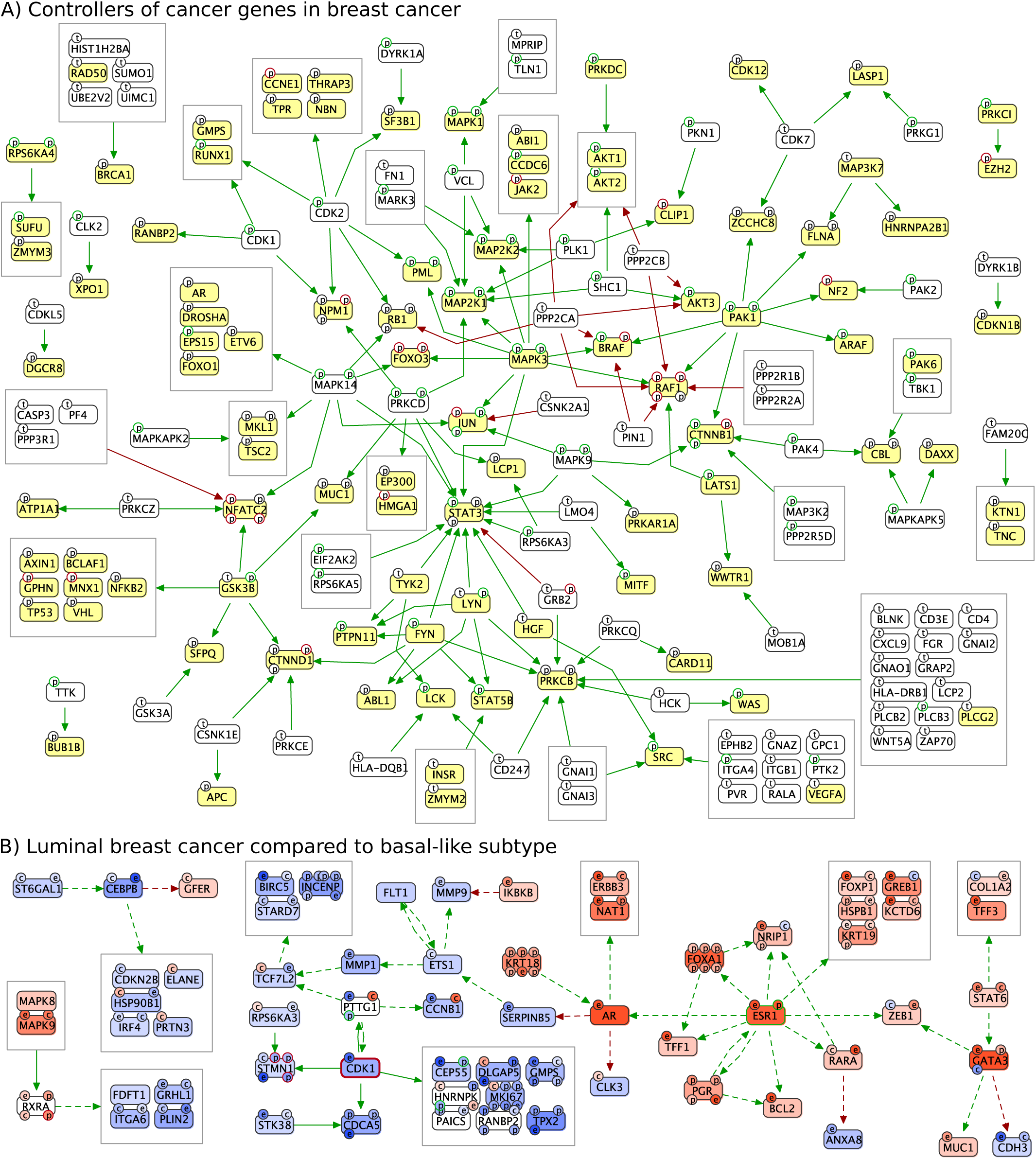
Results for CPTAC breast cancer. A) Subgraph of the correlation-based causal network, showing the upstream regulators of cancer proteins (yellow) in breast cancer. The complete result network without focusing on cancer proteins is given in Supplementary Fig. 3. B) Luminal A and luminal B subtypes are collectively compared to the basal-like subtype. Only the largest connected components are shown. Complete result graph is given in Supplementary Fig. 5.

As a next step in the analysis, we compared the PAM50 expression subtypes of breast cancer to see if we could get causal explanations to their proteomic differences. We were again challenged by decreased sample sizes and missing values, but we detected that both luminal A and luminal B have significant differences from the basal-like subtype. This time, CausalPath results are not significant in terms of the overall network size (*p* = 0.5184), nevertheless, they indicate that ESR1 is significantly more active in luminal breast cancer while CDK1 is significantly more active in basal-like subtype, suggested by both their protein levels and their downstream (Fig. 3B). Transcriptional downstream of ESR1 captures other important transcription factors functioning in the luminal subtypes such as FOXA1, AR and PGR.

### Analysis of TCGA RPPA datasets

There are 32 TCGA projects that provide proteomic and phosphoproteomic measurements with RPPA using over 200 antibodies. The low number of proteins in the datasets prevents a comprehensive pathway analysis, but the antibodies are selected for the proteins’ relevancy to cancer in general and they are typically well studied with many established relations between them. We sought to determine which of these relations most frequently have evidence in the form of correlation across cancer types. We generated a correlation-based causal network for each cancer type using an FDR threshold of 0.001, then we ranked these relations according to how many cancer datasets they can explain (Fig. 4). We found that AKT to GSK3 signaling is the most frequently observed relation, detectable in 30 cancer types, followed by other downstream proteins of AKT including MTOR. Relations between several MAPK signaling proteins, and EGFR to ERBB2 signaling are also among those observed in the vast majority of cancer types. For clarity, here the results do not indicate that these signaling paths are almost always active in cancers, but they indicate that there is almost always a variation in their activity. This is consistent with many studies reporting AKT pathway as a major resistance mechanism to chemotherapy and some other targeted therapies^21–23^.

**Figure 4.**
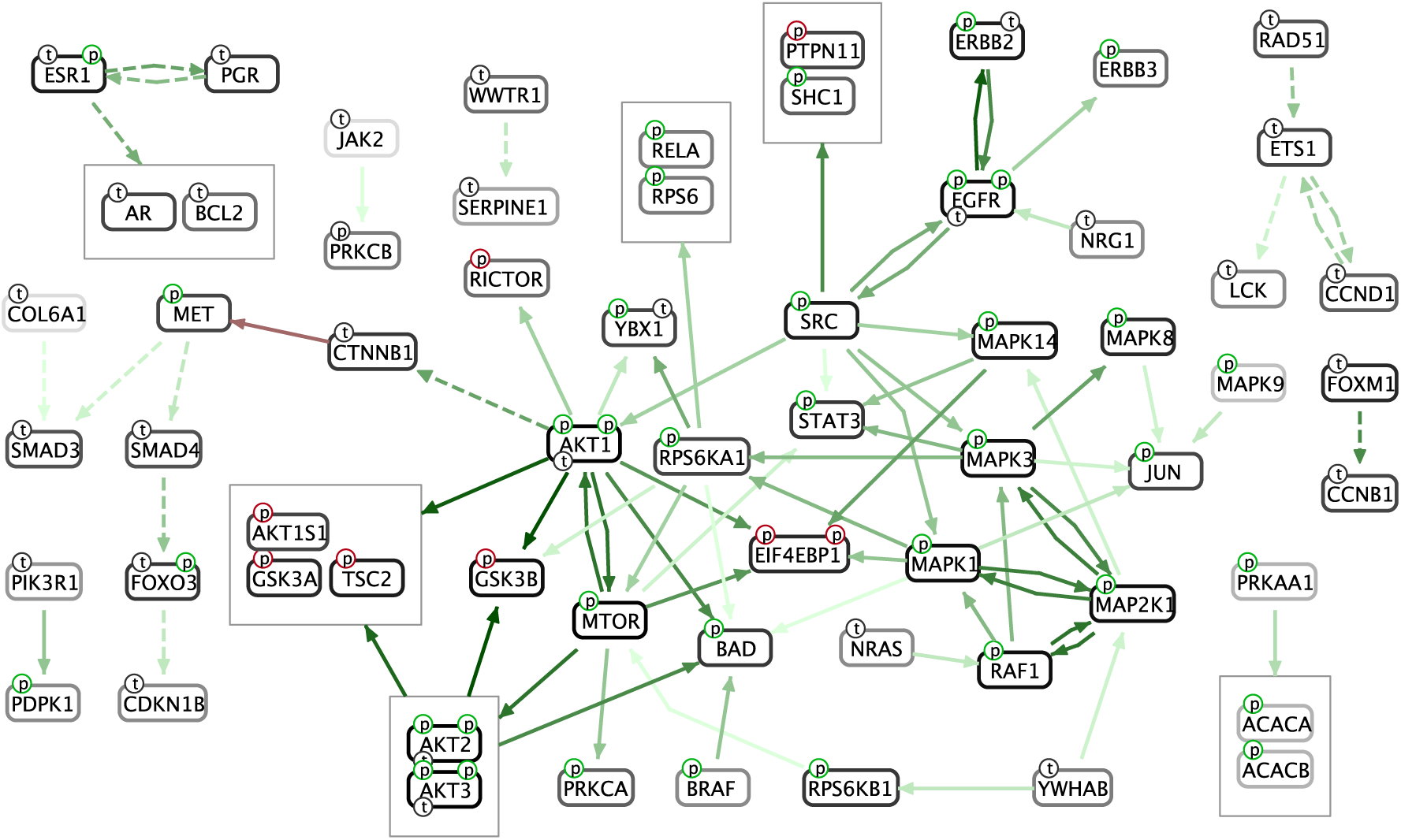
Recurrent results for TCGA RPPA datasets. Relations that are identified with correlation-based analysis in at least 15 cancer types are shown, where faintest color indicates 15 and boldest color indicates 30.

## Discussion

### Pathway extraction versus pathway inference

CausalPath is a novel pathway extraction method to aid researchers in understanding experimental observations using known mechanisms with a focus on post-translational modifications. Experimental data reveal protein features that change in coordination, and CausalPath automates the search for causal explanations in the literature. In other words, correlations come from the data and causality comes from the literature. Compared to the methods that infer causality from data through mathematical modeling (pathway inference), this method has a much wider application area. Pathway inference methods has a potential to offer more accurate results, but they are very data-hungry, requiring many perturbations and/or time points in the experiments, whereas our pathway extraction method is applicable to any simple comparison, or a set of profiles from a cohort with some variance to explain. Additionally, CausalPath is a perfect source for high-confidence *priors*, which can inform the pathway inference methods that benefit from prior data.

### Pathway extraction and biological context

Pathway relations come from experiments performed in highly varied contexts, such as a tissue type, a cell line, gender, or a disease. These relations’ applicability to other contexts is largely unknown, and we are not capable of testing every relation in every context. The approach we present here flattens the pathway data by ignoring the context, then uses proteomic data to select a portion of relations that can explain it. In a way, we let the proteomic data define the context.

### Proteomic data and expressional controls

Our results show that known phosphorylation controls are more consistently observed in the proteomic data compared to known expressional controls. In the ovarian and breast cancer datasets, the size of the resulting phosphorylation networks dramatically decreases with data label shuffling while expression networks remains similar. In the recurrence study with TCGA RPPA datasets, 34% of the resulting phosphorylation controls recur in at least 15 cancer types. This ratio is only 7% for expressional controls. A phosphorylation relation can directly explain a phosphopeptide change, while an expressional control cannot directly explain a total protein change. The latter requires an mRNA change of the target that needs to sequentially cause the proteomic change. While we observe that mRNA and total protein measurements of the same gene are correlated in general (Supplementary Fig. 6), they are obviously not correlated enough to use protein data as a reliable proxy for mRNA. On the other hand, we could identify ESR1 differential activity in breast cancers purely from proteomic data, using expressional relations. Based on these observations, we decided to make it an option in CausalPath, and let users select the molecular data type to use for targets of expressional relations.

### Missing or flawed pathway data

One major limiting factor in this analysis is the large number of protein phosphorylation sites whose functions are not known, hence, their downstream cannot be included in the causality network. We are actively working to mine this data from literature using natural language processing tools, but in the meantime, CausalPath reports those sites with unknown effect which also have significant change at their signaling downstream. Users have the option to review this list of modification sites and manually curate them to increase the coverage of the analysis.

Rarely, in the causality analysis results, we encounter relations that are wrong. These are generally results of faulty manual curation from literature. In these cases, we report them to the source databases, and we remove these erroneous pathway interactions from our network so that future analyses are not affected. We encourage researchers to report such errors to source databases (or alternatively to us) if they come across any to improve the accuracy of our collective knowledge on biochemical pathways.

## Methods

### Significance for proteomic data change and correlation

For comparison-based analyses we used a two-tailed t-test for calculating the significance of the difference of the means of the two groups in the comparison, requiring presence of at least 3 non-missing values from all compared groups. For correlation-based analyses we used Pearson correlation coefficient and its associated significance, requiring at least 5 samples in the calculation. Both tests assume a null model where molecular readouts change independently. The EGF stimulation dataset provides pre-calculated p-values for all of the pairs of time points, which we directly used in the analysis. In all calculations, we used the Benjamini-Hochberg method for controlling FDR, whenever applicable. We used 0.1 as a default FDR threshold, unless indicated otherwise.

### Causality

Both pathway inference and pathway extraction are closely related to formal notions of causality inference, specifically to the Suppes’ and Pearl’s probabilistic formulations. A probabilistic causal relationship between two events, say from event *A* to event *B*, indicates the probability of *B* depends on the status of *A*, as described by Patrick Suppes^24^. While using this notion of causality can generate predictive models, it does not tell if *A* may cause *B*. For instance, there can be an event *X* that is causing both *A* and *B*, and this will still satisfy Suppes’ formulation. To make the model predictive under an intervention scenario, Judea Pearl provided a reformulation: perturbing the status of *A* will change the probability of *B*^25^. We follow Pearl’s notion and detect mechanism-based evidence for activity change of one protein may affect abundance of a specific peptide from another protein in pathway databases, as described in the next section.

### Derivation of binary relations from detailed mechanistic pathway data

Using the BioPAX-pattern framework, and by studying the structure of the BioPAX models from different resources, we manually defined 12 BioPAX patterns to capture potentially causal binary relations that involve phosphorylation and expression of proteins. We provide the details of these patterns in the Supplementary Document, along with examples for what they can detect.

### Description of causal graph notation

We developed a new graph notation to represent resulting causal explanations as a logical network, where nodes denote proteins and edges denote causal relations (Fig. 2C). Node background is used for color-coding total protein change, while site-specific changes are shown with small circles on the nodes whose border colors indicate whether the site is activating/inhibiting. If additional omic data such as RNA expression, DNA copy number, or mutation status are available, we include them for integrated visualization, using small circles displaying specialized letters. Binary causal relations are represented with edges—green representing positive, red representing negative, phosphorylations with solid edges and transcriptional regulations with dashed edges. When significance calculation results are available, they are represented on the protein borders, using a bold border when downstream of a protein is significantly large, green border when downstream indicates the protein is activated, and red border when downstream indicates the protein is inactivated. If both the activation-indicating and inhibition-indicating downstreams are significantly large, then a dark yellow color is used instead. We use topology grouping while rendering result networks, which means we group the proteins with the same network topology under compound nodes on the network for complexity management.

### CausalPath parameters

CausalPath is designed to explore omic datasets of different sizes, types and accuracy. Following are some important parameters to consider when using the method.

#### Site matching proximity threshold

Protein phosphorylation sites in the literature have to exactly match the detected site in the phosphoproteomic dataset to use in causal reasoning by default. Some users may find this too strict since there can be slight shifts in the literature, or some nearby sites of proteins are likely to be phosphorylated by the same kinase. This parameter lets the analysis allow a determined inaccuracy in site mapping to explore such cases. We used strict site-matching for all the CausalPath analyses unless indicated otherwise.

#### Site effect proximity threshold

The effect of the phosphorylation sites on the protein activity, as in activating or inactivating, are curated by pathway databases, mostly by PhosphoSitePlus. We also did some small scale curation for platelet analysis, EGF stimulation analysis, and RPPA analyses. CausalPath requires exact matching of these curated site effects with the sites in the data by default, however this can be too strict because nearby sites generally tend to have similar effects. This parameter lets the analysis use a determined inaccuracy while looking up site effects. In this manuscript we always used accurate matching for site effects.

#### Protein focus

This parameter lets the analysis use a subset of the literature relations, focusing on the neighborhood of certain proteins indicated by their gene symbols, hence reducing the number of tested hypotheses. We used this parameter during analysis of ovarian cancer subtypes as described in the relevant section.

#### Generation of a data-centric causal network

CausalPath result networks are gene-centric, meaning that genes are represented with nodes, and all other measurements related to a gene are mapped on the gene’s node. When a data row can map to multiple genes, however, this creates a redundancy, as we have in the RPPA analysis results. For example, the AKT phosphoantibody can recognize all three AKTs, so we duplicated the same data on AKT1, AKT2 and AKT3. A similar problem exist for mass spectroscopy when a phosphopeptide can be resolved to multiple orthologous proteins, if their sequences are identical around the phosphorylation site, AKTs again being an example. As an alternative to this view, CausalPath can generate data-centric views where nodes represent data rows unresolved to particular proteins. But this view do not support mapping other available -omic data onto the network, and also the relations are duplicated when more than one data of the same gene can be explained by the same relation.

#### Protein activity

CausalPath allows users to insert their own hypotheses about protein activity changes in the case of a comparison-based analysis. The input has to be a gene symbol associated with a Boolean parameter indicating the hypothesized direction of activity change, i.e. activated or inhibited. We used this option for EGF stimulation analysis to indicate that we expect EGF to be activated because the data is only phosphoprotein, hence there is no measurable change on EGF itself to include in the analysis otherwise.

#### Data type for expressional targets

CausalPath keeps its logical reasoning within proteomic data by default, however users can opt to use mRNA data for the targets of expressional relations. We used this parameter during correlation-based analysis of ovarian and breast cancers as described in the results section. It is also possible to use mRNA and protein data together by using this parameter multiple times.

### Robustness

We tested the robustness of CausalPath against noise on the CPTAC breast cancer dataset, on a correlation-based analysis with two different FDR thresholds. We iteratively introduced noise into the dataset by adding random numbers drawn from a normal distribution, and generated the causal network. At each step, we recorded the amount of the original relations as well as new relations that we have in the results (Fig. 5). Ideally, as the noise level increases, we would like the method to retain the original results and not allow the noise to generate new relations. Our tests indicate that, the original relations are sensitive to noise, which means they rapidly start decreasing in number, however, the method is safe against noise, which means we do not get many new relations due to it.

**Figure 5.**
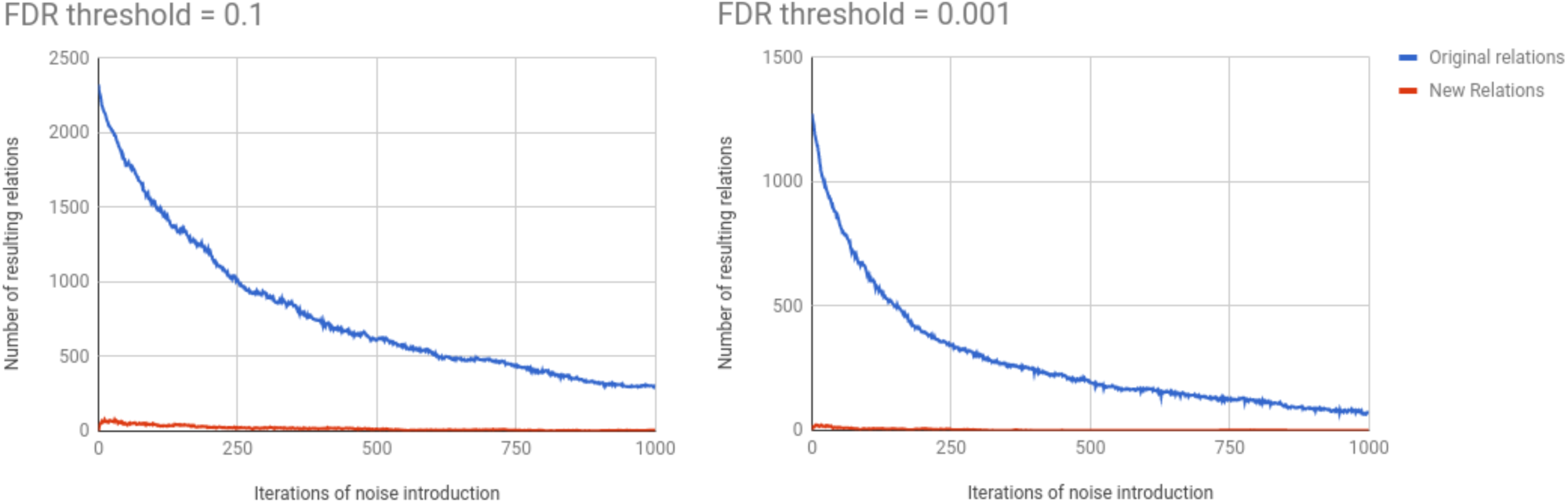
Robustness of CausalPath on CPTAC breast cancer dataset.

### Reproducibility

We tested the reproducibility of CausalPath on the CPTAC breast cancer dataset by using random subsets of the samples, on a correlation-based analysis with two different FDR thresholds. In total of 100 trials, we used a random half of the samples at each step, and checked how frequently each causal relation is reproduced in the results (Fig. 6). The results indicate that 70% of the original relations are reproduced consistently in at least half of the trials when FDR threshold is 0.1. This ratio is reduced to 49% when we use an FDR threshold of 0.001. The significant amount of new relations (red) with low reproduction counts in the tests are due to the accumulation of false positives from all 100 trials.

**Figure 6.**
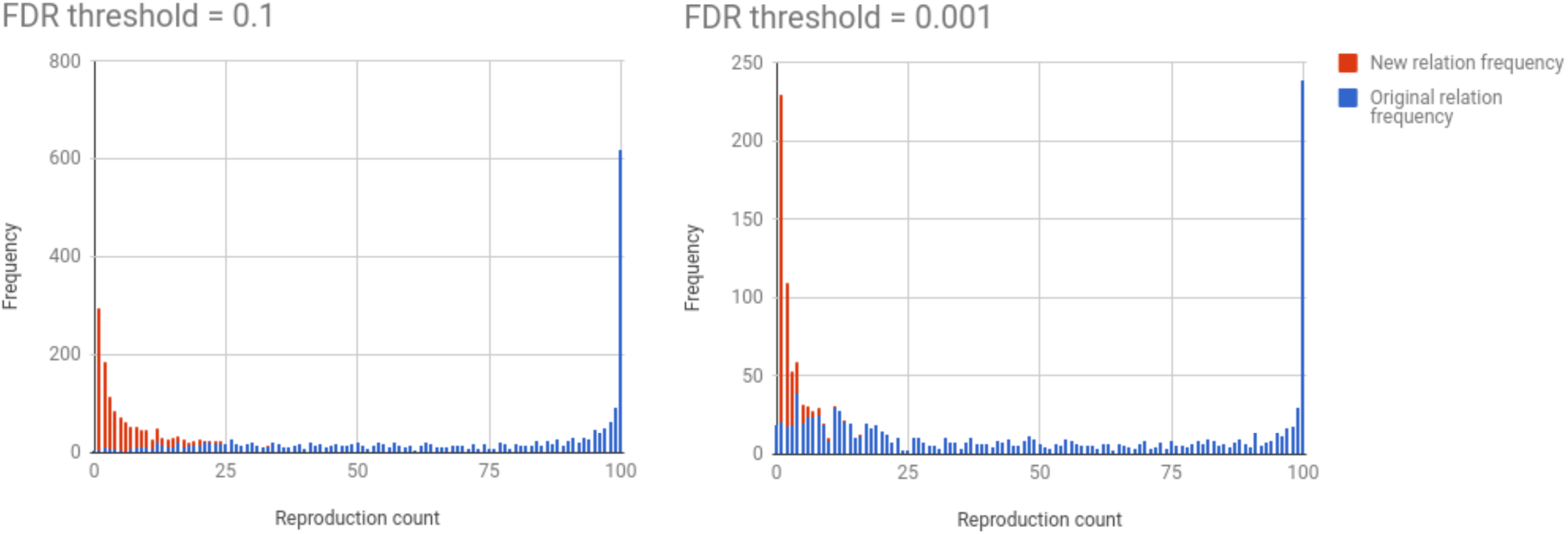
Reproducibility of CausalPath on CPTAC breast cancer dataset.

### Previously published methods for pathway analysis of proteomic datasets

There is no method comparable to CausalPath for its ability to identify causal relations from pathway databases that can explain given proteomic datasets. But there are methods developed for other forms of pathway analysis for proteomics. Below is a short survey of these methods.

#### pCHIPS^26^

This is a network propagation method for proteomic and other data, based on the TieDIE^27^ algorithm, where the purpose is to link differentially active kinases (indicated by proteomic data) to the differentially active transcription factors (indicated by RNAseq measurements of targets). Proteomic changes on the kinases are propagated downstream, differential transcription factor activities are propagated upstream on the signaling network, and the overlap is identified as possible linking path(s).

While linking kinases to the transcription factors implies causality, pCHIPS does not check the conditions of causality such as if the proteomic change is indicative of activation or inhibition, or if the linking path has a positive or a negative effect on the transcription factor activity, or their compatibility for a causality hypothesis.

#### PhosphoPath^28^ and PTMapper^29^

Both methods are implemented as a Cytoscape plugin to visualize kinase-substrate relations on the protein-protein interaction (PPI) network. Users can run a network enrichment analysis on the PPI network for the given proteomic and other datasets, then visualize the known kinase-substrate relations on the enriched region.

#### PCST^30^

This method maps proteomic and transriptomic data on the proteins on a weighted PPI and protein-DNA interaction network, then identifies a minimal subnetwork that connects the mapped molecules, prioritizing the most reliable connections. Authors formulate this as a prize-collecting Steiner tree (PCST) problem, and solve with a known algorithm.

#### PHOTON^31^

This method maps proteomic data from a perturbation study onto the proteins on a weighted PPI network, then calculates a score for each protein based on the weighted average of the observed proteomic changes on its neighbors on the network The method generates a result network by connecting the perturbed protein and the proteins with high score on the PPI network.

### Availability

CausalPath is freely available at http://causalpath.org. For an analysis, users need to provide the proteomic data in a special tab-delimited format, along with the analysis parameters such as how a “change” should be derived from the data. Options include averaging a group of values, getting difference/fold-change of two groups of columns, compare two groups with a t-test, or use correlations in a single group. The results are displayed on the browser with the utilization of Cytoscape.js^32^, and the mechanistic details of the causal interactions can be viewed in SBGN-PD language, which is implemented by the utilization of SBGNViz^33^. An alternative way to run CausalPath for computational biologists is from its Java sources, which is freely available at https://github.com/PathwayAndDataAnalysis/causalpath. The generated result networks can either be viewed by uploading to the web server or by loading into ChiBE. All network figures in this manuscript are generated with ChiBE.

## Acknowledgements

This work was sponsored by DARPA under the Big Mechanism Program (Contract # W911NF-14-C-0119) and the U. S. Army Research Office (Contract # ACC-APG_RTP W911NF), and by NIH grants U41HG006623 (Pathway Commons) and P41GM103504 (National Resource for Network Biology). Anil Korkut is supported by MD Anderson Cancer Center Support Grant P30 CA016672 (Bioinformatics Shared Resource). We thank to Hannah Manning and Olga Nikolova for critical reading of the manuscript. The results published here are in part based upon data generated by the TCGA Research Network: http://cancergenome.nih.gov/.

